# Adaptive diversification of growth allometry in the plant *Arabidopsis thaliana*

**DOI:** 10.1101/269498

**Authors:** François Vasseur, Moises Exposito-Alonso, Oscar Ayala-Garay, George Wang, Brian J. Enquist, Denis Vile, Cyrille Violle, Detlef Weigel

## Abstract

Seed plants vary tremendously in size and morphology. However, variation and covariation between plant traits may at least in part be governed by universal biophysical laws and biological constants. Metabolic Scaling Theory (MST) posits that whole-organismal metabolism and growth rate are under stabilizing selection that minimizes the scaling of hydrodynamic resistance and maximizes the scaling of resource uptake. This constrains variation in physiological traits and in the rate of biomass accumulation, so that they can be expressed as mathematical functions of plant size with near constant allometric scaling exponents across species. However, observed variation in scaling exponents questions the evolutionary drivers and the universality of allometric equations. We have measured growth scaling and fitness traits of 451 *Arabidopsis thaliana* accessions with sequenced genomes. Variation among accessions around the scaling exponent predicted by MST correlated with relative growth rate, seed production and stress resistance. Genomic analyses indicate that growth allometry is affected by many genes associated with local climate and abiotic stress response. The gene with the strongest effect, *PUB4*, has molecular signatures of balancing selection, suggesting that intraspecific variation in growth scaling is maintained by opposing selection on the trade-off between seed production and abiotic stress resistance. Our findings support a core MST prediction and suggest that variation in allometry contributes to local adaptation to contrasting environments. Our results help reconcile past debates on the origin of allometric scaling in biology, and begin to link adaptive variation in allometric scaling to specific genes.

**Significance statement:** Are there biological constants unifying phenotypic diversity across scales? Metabolic Scaling Theory (MST) predicts mathematical regularity and constancy in the allometric scaling of growth rate with body size across species. Here, we show that adaptation to climate in *Arabidopsis thaliana* is associated with local strains that substantially deviate from the values predicted by MST. This deviation can be linked to increased stress tolerance at the expense of seed production, and it occurs through selection on genes that are involved in abiotic stress response and that are geographically correlated with climatic conditions. This highlights the evolutionary role of allometric diversification and helps establish the physiological bases of plant adaptation to contrasting environments.

## Introduction

At the core of the quest for understanding and predicting biological diversity is the apparent paradox that, despite the phenotypic changes that underlie divergent ecological strategies, there seem to be constant or near-constant parameters across life forms (1). The latter is assumed to result in part from biophysical constraints limiting the range of possible trait values (2), as well as from strong stabilizing selection for optimal phenotypes (3, 4). Consistently, body size variation in multicellular organisms is associated with many scaling regularities. Max Kleiber (5) first reported that the consumption of energy (metabolic rate *G*) varies to the ¾-power of organism mass *M*, such that *G* = *G*_*0*_*M*^*3/4*^, implying that a 10-fold increase in *M* produces in virtually all organisms a 5.6-fold increase in *G*. Several physiological models have been proposed to explain this constancy. The most prominent is Metabolic Scaling Theory (MST) (6), which predicts that scaling exponents of several traits tend to take on “quarter-power” values (*e.g.*, ¾, ¼) as the outcome of an optimal balance between the scaling of hydraulic transport costs and the scaling of exchange surface areas (*e.g.*, leaf area in plants) (7). According to MST, the scaling of physiological rates matches the ability of exchange surfaces to obtain resources from the environment and then distribute them to metabolizing cells through the vascular network. Because the branching geometry of this network is highly constrained in space, it is predicted that selection that minimizes the costs of resource transport and at the same time maximizes the uptake of resources will lead to “allometrically ideal” organisms characterized by a common set of quarter-power scaling relationships with body mass.

Empirical observations support MST predictions across land plants, where several traits, including organismal growth rate, scale as body mass raised to the power of ¾ (8, 9). On the other hand, the scaling exponent can vary across plants (10–12), or scaling can be constant but deviate from ¾ (13). These seemingly contradictory observations have been proposed to reflect (i) phenotypic, like life history, differences between species or populations (9, 10), (ii) physiological changes along environmental gradients (14, 15), or (iii) non-linearity in hydrodynamic resistance and metabolic scaling (16). Thus, important questions about the evolution of allometry remain (4). For example, is the prevalence of ubiquitous scaling relationships the result of stabilizing selection acting to remove unfit genetic allometric variants? And does variation in the scaling exponent reflect adaptation and genetic diversification, or developmental plasticity?

To address these and related questions, we examined how growth rate scales with body size in a genetically diverse population of *Arabidopsis thaliana* accessions (Dataset S1), a species that exhibits three orders of magnitude in plant dry mass (10) and occurs in a wide range of contrasting environments (17). We provide evidences that scaling variation is maintained by an adaptive trade-off between alternative environments. We show that this variation has a polygenic basis, and that there is genetic correlation between allometry and local climate.

## Results

### Variation of *A. thaliana* Growth Scaling with Climate

The scaling exponent of growth is conventionally quantified as the slope θ of the allometric function y = α + θx, where x and y are the logarithms of plant biomass and absolute growth rate, respectively. Fitting the allometry of the mean absolute growth rate (GR, mg d^−1^), estimated as the ratio of final plant dry mass (mg) over total duration of the life cycle (days), across *A. thaliana* accessions returned a scaling exponent θ that is not significantly different from the MST predicted value of ¾ (y = −1.07 + 0.74x; *r*^2^ = 0.97; slope CI_95%_ = [0.725, 0.750]; Fig. 1A). This value is the same as observed across vascular plant species (box in Fig. 1A). However, the relationship is not a pure power function, and instead was better explained by a non-linear quadratic function (y = −1.93 + 1.43x − 0. 14x^2^, ΔAIC = −192.4; Fig. 1A). Our analyses indicate that this curvilinear scaling relationship was due to differences in θ between accessions, which can be estimated as the first derivative of the quadratic function (θ = 1.43 - 0.27x), and which varied between accessions from 0.47 to 1.10 (Fig. 1B, Fig. S1C). The broad-sense heritability, *H*^2^, of θ was 0.95, which is higher than any other trait measured in this study (Table S1), indicative of a high amount of variance explained by genetic effects in our highly controlled growth conditions.

**Figure 1.**
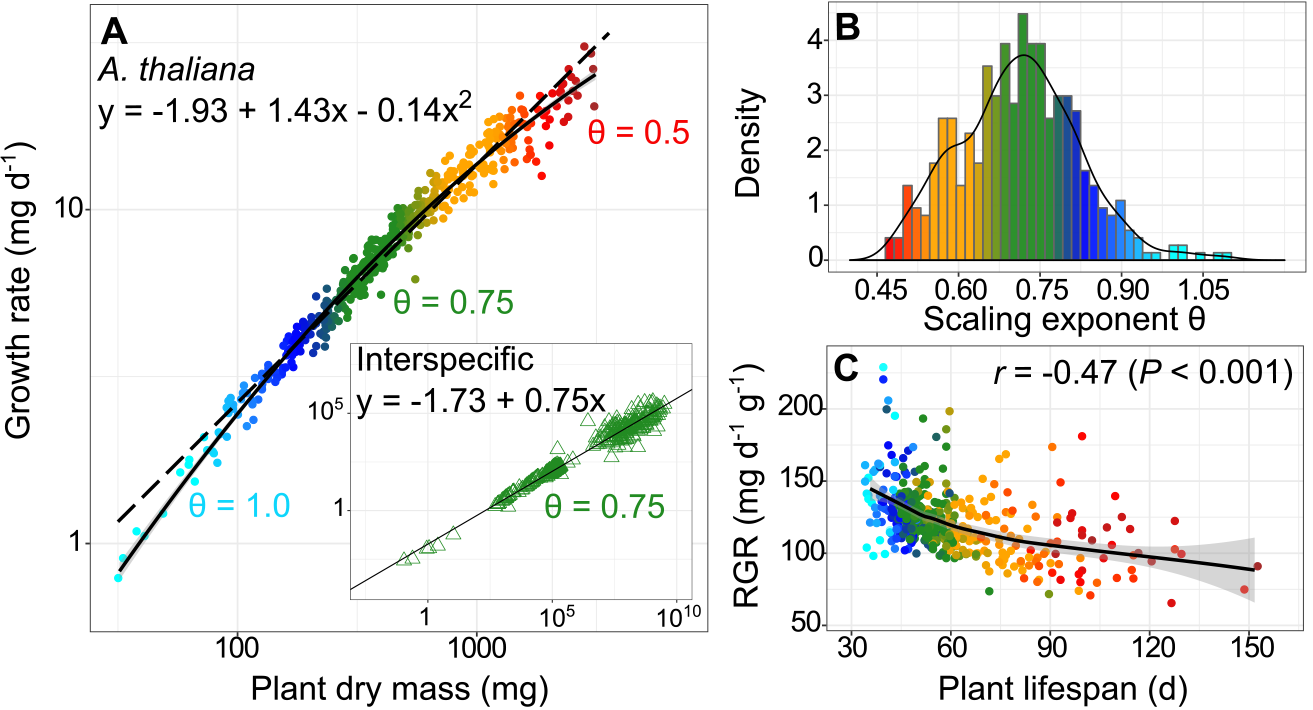
Variation of growth scaling in *A. thaliana*. (**A**) Linear (dashed line) and quadratic (solid line) fits of mean growth rate versus final dry mass in 451 *A. thaliana* accessions. Box: linear fit (black line) of growth rate versus plant dry mass in 333 vascular plant species from Niklas and Enquist (8). (**B**) Distribution of the scaling exponent derived from the quadratic fit in the 451 *A. thaliana* accessions. (**C**) Relationship between relative growth rate (RGR) at growth maximum, plant lifespan and scaling exponent in the 451 accessions. Black curve is Loess fit ± 95% CI (grey area). In all panels, dots and triangles represent genotypic and species means, respectively, colored by the value of the scaling exponent reported in panel (**B**).

Modelling the dynamics of plant dry mass accumulation from imaging data (18) revealed that the estimated relative growth rate (RGR) explains 18% of the variation in the scaling exponent (*P* < 0.001), with both being negatively correlated with plant lifespan (*P* < 0.001, Fig. 1C; Dataset S2). Previous studies have shown that variation in *A. thaliana* growth allometry is positively correlated with carbon assimilation rate and nutrient concentration, but negatively with lifespan (10). Thus, variation of growth allometry in *A. thaliana* connects life-history variation to the strategies for leaf resource-use. At the one end of the distribution are high scaling exponents, representative of ‘live fast/die young’ strategies that maximize resource capture (high RGR and carbon assimilation rate) at the expense of plant lifespan and final size. At the other end are low scaling exponents, representative of ‘live slow/die old’ strategies that maximize the retention (thick leaves with low nutrient concentration and long lifespan) rather than acquisition of resources.

We then examined the correlations between the scaling exponent and 21 climatic variables, which include 19 ‘Bioclim’ variables (http://www.worldclim.org/bioclim), as well as the estimated mean annual Potential Evapo-Transpiration (PET, mm) and Aridity Index (19) at the geographic origin of the accessions. Consistent with the idea that resource-acquisitive plants, *i.e.* early-flowering/fast-growing ecotypes, are more adapted to hotter and drier regions, the scaling exponent was positively correlated with the mean annual temperature measured at the collection point of the accessions (Fig. 2A; Dataset S2). The strongest correlations were with maximum temperature of the warmest month and mean temperature of the warmest quarter (*r* = 0.30 and 0.28, respectively, Fig. 2B; Dataset S2). Inversely, the scaling exponent was negatively correlated with precipitation, specifically with precipitation during the driest quarter (Fig. 2C), precipitation seasonality and the aridity index (Dataset S2). In contrast, it was not correlated with the altitude at the collection point.

**Figure 2.**
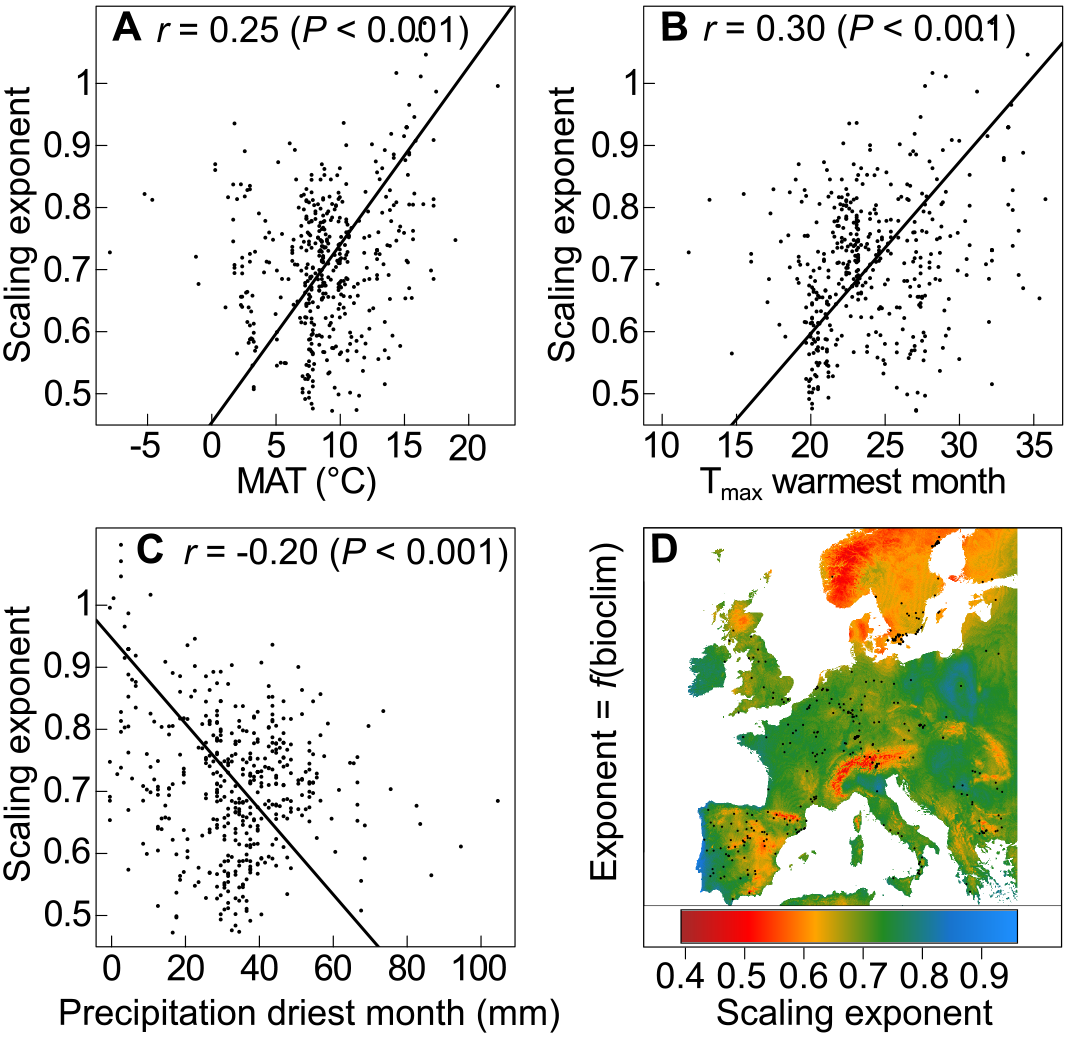
Relationships between scaling exponent and climate. (**A-C**) Correlations between the scaling exponent measured across the 451 accessions and local mean annual temperature (**A**), maximum temperature of the warmest month (**B**), and precipitation of the driest month (**C**). Dots represent genotypic mean. Fitted lines are SMA regressions. *r* is the Pearson’s coefficient of correlation with associated *P*-value. (**D**) Geographic distribution of the scaling exponent across Europe in *A. thaliana*, modelled as a function of 13 Bioclim variables. Colors indicate the predicted value of the scaling exponent. Black dots represent geographic origins of the accessions phenotyped.

Using stepwise regression, we found that 13 climatic variables explain >27% of the allometric variation. Four of these are related to summer and two to winter climate. The strongest effects were estimated for annual mean temperature, isothermality and mean summer temperature. Modeling the geographic distribution of scaling exponent with the 13 top-correlated climatic variables as predictors showed that intermediate exponents are more common in temperate regions (Fig. 2D), while extreme exponents are favored under more stressful conditions (*e.g.* high altitude, high latitude).

### Fitness Costs and Benefits of Allometric Variation

The scaling exponent was correlated with resource-use traits including RGR and lifespan, as well as performance-related traits such as fruit number, a proxy for lifetime fitness in annual species (fruit number varied from 18 to 336 per plant, Table S1; *SI Appendix*). However, the relationship between fitness and the scaling exponent under the non-limiting RAPA conditions was not linear (Fig. 3A). Instead, fruit number was a bell-shaped function of the scaling exponent: it peaked for plants with an exponent around ¾ and declined towards higher or lower exponents. Thus, genetic deviations from the ¾ scaling exponent are associated in *A. thaliana* with extreme resource-use strategies, and a general decline in fruit number (*r* = −0.62, *P* < 0.001; Dataset S2). A polynomial regression of relative fitness - using fruit number standardized by the population mean - over the scaling exponent returned a significant, negative second-order coefficient (y = 1.00 +4.23x − 4.06x^2^, *P* < 0.001 for all coefficients), *i.e.* an estimate of quadratic selection gradient |γ| that might be indicative of stabilizing selection for the allometric exponent under benign conditions (20).

**Figure 3.**
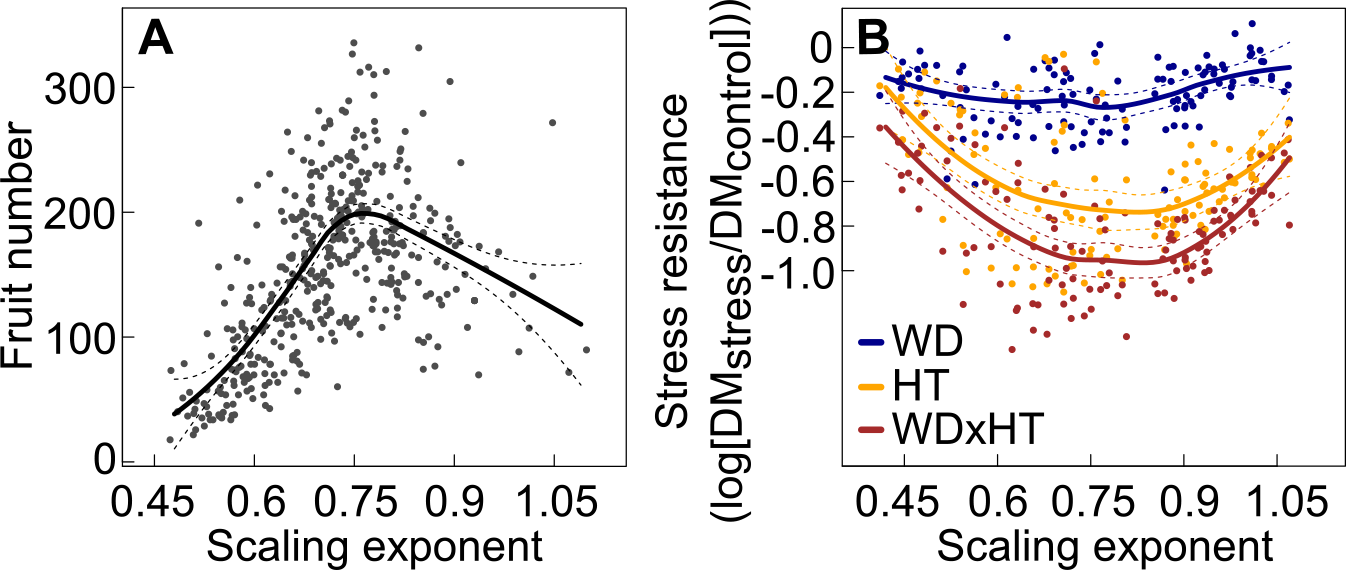
Relationships between scaling exponent, fitness and resistance to abiotic stress. (**A**) Relationship between fruit production and scaling exponent in the 451 accessions. Black curve is Loess fit ± 95% CI (dashed lines). (**B**) Stress resistance expressed as the log_10_ of the ratio of final rosette dry mass under water deficit, high temperature, and both compared to control conditions, across 120 *A. thaliana* recombinant inbred lines. Data have been published (10, 23). Dots indicate genotypic means (*n* = 4). Colored curves are Loess fit ± 95% CI (dashed lines).

Conversely, deviation from ¾ scaling was positively correlated with survival under severe drought (*r* = 0.16, *P* < 0.05; measured in (21) across 210 common accessions; Dataset S2), and negatively correlated with growth reduction under moderate drought (*r* = −0.26, *P* < 0.05; measured in (22) across 60 common accessions, Dataset S2). However, neither stress-resistance trait was correlated with the scaling exponent itself. This suggests that deviation of allometric exponents from ¾ in either direction is associated with increased resistance to stressful conditions at the expense of reduced reproductive fitness under benign conditions. Consistently, a re-analysis of an experimental population phenotyped for tolerance to combined high temperature and water deficit (23) pointed to higher stress sensitivity of accessions with scaling exponents close to ¾ (Fig. 3B). In contrast, allometric exponents at both the low and high end of the distribution were correlated with improved stress tolerance, specifically under high temperature (Fig. 3B). A possible explanation of this result could be that a ‘fast’ strategy with high scaling exponents allows stress escape by maximizing resource acquisition and completion of the life cycle before a short window of non-stressful conditions closes (23). Alternatively, the ‘slow’ strategy might support stress tolerance by reducing metabolic activities and thus, the resource demand associated with a fast growth (10).

### The Genetic and Evolutionary Bases of Allometric Variation

Because we suspected that allometric variation might result from adaptation to the diverse environments at the places of origin of accessions, we looked for genetic evidence of local adaptation and of genetic diversification with climate. Principal component analysis (PCA) performed after eigen decomposition of the relatedness matrix revealed that the scaling exponent was correlated with population structure, notably with the second PCA axis (*r* = 0.37, *P* < 0.001), which explains 28% of total genetic variation and mainly differentiates accessions from Relicts, N. Sweden and Spain groups (17) (Fig. S2). By contrast, flowering time was correlated with the first PCA axis, which explains 42% of genetic variation and is associated with longitudinal divergence among accessions (Fig. S2). Compared to the ancestral (‘Relict’) genetic group (17), scaling exponent differed significantly (*P* < 0.001) for two groups: N. Sweden and S. Sweden, while the eight other groups were not different (*P* > 0.3). Q_st_ of scaling exponent - measured as the ratio of between-group phenotypic variance over total variance - was above 0.9 quantile of genome-wide F_st_ (Q_st_/F_st_ ratio = 2.14, *P* < 0.001; Table S1, Fig. S3), which is potentially indicative of polygenic selection acting on the scaling of plant growth (24).

We ran GWA models on the scaling exponent θ and the 21 climatic variables using the EMMAX procedure to correct for population structure (25). In total, 8,250 single nucleotide polymorphisms (SNPs) out of 1,793,606 tested were significantly associated with at least one phenotypic trait or climatic variable (Dataset S3) after multiple-testing correction (26). Only six SNPs were significantly associated with the scaling exponent (FDR < 0.05). Five of these six SNPs were located in the same region on chromosome 2 (Fig. 4A), and were associated with maximum temperature of the warmest month (Fig. 4B). Three SNPs were also significantly associated with the mean annual temperature and the mean temperature of the coldest month (Dataset S4). The same genomic region showed strong association with precipitation during the driest month (Fig. 4C), although the six SNPs that were associated with scaling variation did not reach the significance threshold for this climatic variable (FDR > 0.05). In contrast, no significant SNPs were shared between RGR, lifespan, fruit number or rosette dry mass and the climatic variables (Dataset S4), suggesting that genetic association between traits and climate is relatively rare.

**Figure 4.**
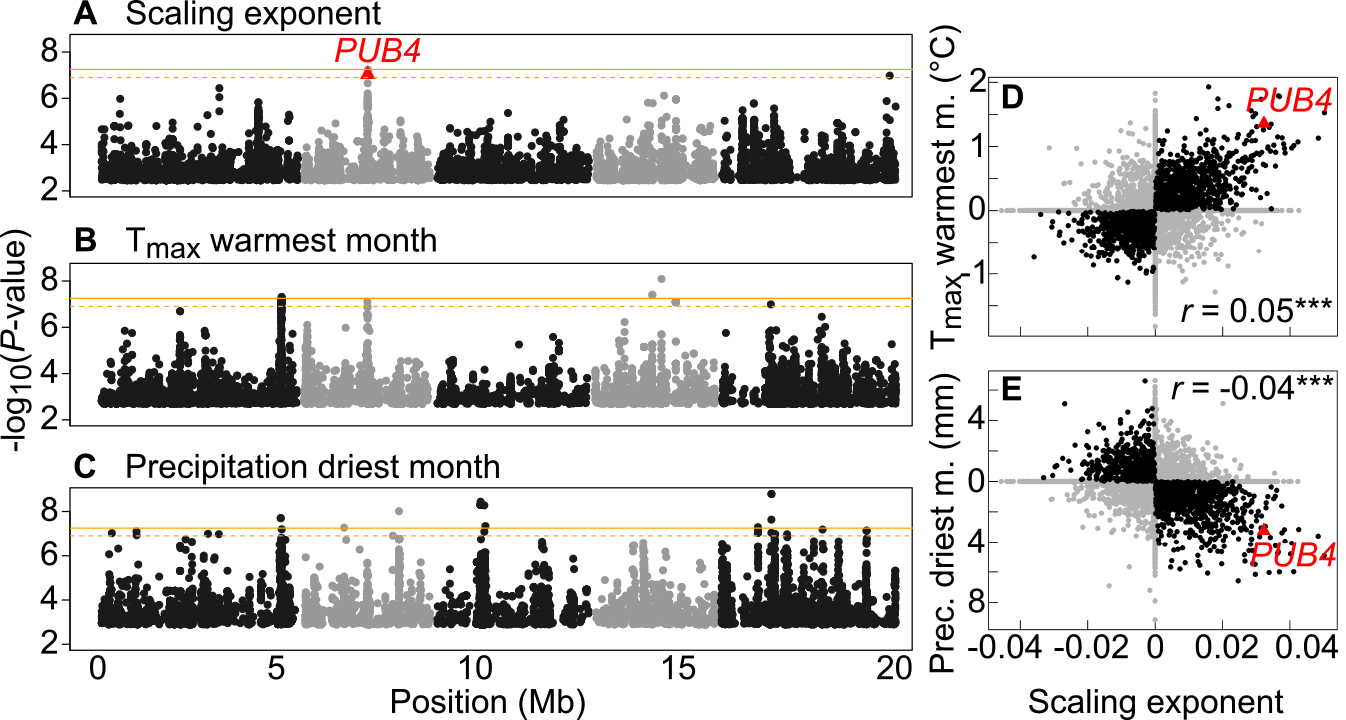
GWA mapping of allometric variation in *A. thaliana*. (**A-C**) Test statistics for SNP associations (EMMAX) with (**A**) scaling exponent, (**B**) maximum temperature during the warmest month, and (**C**) precipitation during the driest month. Dots are 1% top-associated SNPs along the five chromosomes (alternate grey and black dots represent chromosomes). Orange lines represent genome-wide significance threshold with Bonferroni correction at α = 0.05 (solid line) and α = 0.1 (dashed line). Red triangle is *PUB4* (FDR < 0.05) (**D**, **E**) Correlation between SNP effects (BSLMM) for scaling exponent and maximum temperature of the warmest month (**D**), and precipitation of the driest month (**E**). Black dots represent similar SNP effect for x and y variables (both positive or both negative). *r* is Pearson’s coefficient of correlation (***: *P* < 0.001).

One SNP among the five associated with both the scaling exponent and the maximum temperature of the warmest month was located in the U-box protein gene *PUB4* (At2g23140; MAF = 6.1%; Fig. 4A). As E3 ubiquitin ligases, U-box proteins are involved in protein turnover, a key regulatory component of plant responses to abiotic stresses (27). PUB4 plays notably a role in a quality-control pathway that removes damaged chloroplasts (28). Two other SNPs were located in the nearby cytochrome P450 gene *CYP81D6* (At2g23220), 40 kb from *PUB4* (*r*^2^ = 0.63). CYP450s catalyze the production of diverse secondary metabolites that are involved in biotic and abiotic stress response (29). The remaining two SNPs were also linked to *PUB4* and *CYP81D6*, but affected non-coding sequences. We note that the *PUB4* polymorphisms only account for about 1% of the genetic variance in the scaling exponent. Because broad-sense heritability was *H*^2^ > 95%, many other loci are expected to contribute to allometric variation, potentially reducing the power of classical GWA to detect SNPs significantly associated with the scaling exponent. For instance, we expected that, given the strong correlation between the scaling exponent and plant lifespan (Dataset S2), many flowering time genes would be significantly associated with allometry. However, no SNP reached the significance threshold for lifespan in our analysis (FDR > 0.05), and we therefore do not have evidence for flowering time genes being predictors of allometric variation. This might be due to over-correcting for population structure, or to the high number of SNPs involved in phenotypic variation between accessions. Indeed, a strong correction for population structure might be inappropriate if many genes across the entire genome contribute to the phenotype in question.

To account for the potentially complex genetic architecture of traits, we ran Bayesian Sparse Linear Mixed Models (BSLMM) implemented in GEMMA (30). BSLMM models two hyperparameters, a basal effect α_i_ that captures the fact that many SNPs contribute to the phenotype, and an extra effect β_i_ that captures the fact that not all SNPs contribute equally. SNP effects, which can be estimated as the sum of α_i_ and β_i_ (30), were strongly correlated between the scaling exponent and all climatic variables except temperature annual range (Dataset S5). As expected, correlations between SNP effects on scaling exponent and climate were strongest for mean annual temperature, and temperature and precipitation during summer (Dataset S5). Consistent with the measurement of broad-sense heritability (*H*^2^), ‘chip’ heritability - a proxy for narrow-sense heritability (*h*^2^) measured with GWA - was very high for the scaling exponent (*h*^2^ = 0.87 versus *H*^2^ = 0.95; Table S1), suggesting that most of the phenotypic variance can be explained by the additive effects of SNPs controlling allometric variation.

Gene ontology (GO) analysis (31) of the 1% top-genes affecting the scaling exponent revealed enrichment in genes with catalytic activity and ones related to carbohydrate metabolism, post-embryonic development, post-translational protein modification, and response to abiotic stimulus (Fig. S4A, B). A large fraction of the proteins encoded by these genes are predicted to localize to plasma membranes or the chloroplast (Fig. S4C). F_st_ values across the 100 top-genes were significantly higher than genome-wide F_st_ values (F_st [100 top-genes]_ = 0.23 versus F_st [Genome-wide]_ = 0.17, *P* < 0.001; Fig. S3), which is consistent with Q_st_ analysis and indicative of polygenic selection on the genes controlling growth allometry. As expected, *PUB4* is among the 100 top-genes associated with plant allometry, showing strong effects on both the scaling exponent and climatic variables (Fig. 4D, E). We estimated that *PUB4* alone favors plant adaptation to warmer and drier summers by up to +1.4 °C and −3mm (Fig. 4D, E) through an increase of the scaling exponent by up to +0.03.

A scan for genomic signatures of selection in the 50 kb region around *PUB4* revealed increased Tajima’s D (Fig. 5A) and SNP-level F_st_ (Fig. 5B), but we did not observe signatures of recent selection sweeps. As an index of allelic diversity that quantifies departures from the standard neutral model (32), high Tajima’s D values indicate an excess of intermediate-frequency alleles, a potential sign for balancing selection, specifically in *A. thaliana* where Tajima’s D is commonly negative due to recent population expansion and selfing (33, 34). This is consistent with molecular signatures of climate selection previously observed in *A. thaliana* (35, 36). Moreover, climate-envelope modelling of *PUB4* allelic distribution revealed strong geographic structure associated with summer conditions; the major *PUB4* allele is mostly found in temperate and cold northern parts of Europe (Fig. 5C), while the minor allele is mostly Mediterranean (Fig. 5D). This supports the role of *PUB4* in evolutionary adaptation to warmer and drier regions around the Mediterranean through variation in growth scaling.

**Figure 5.**
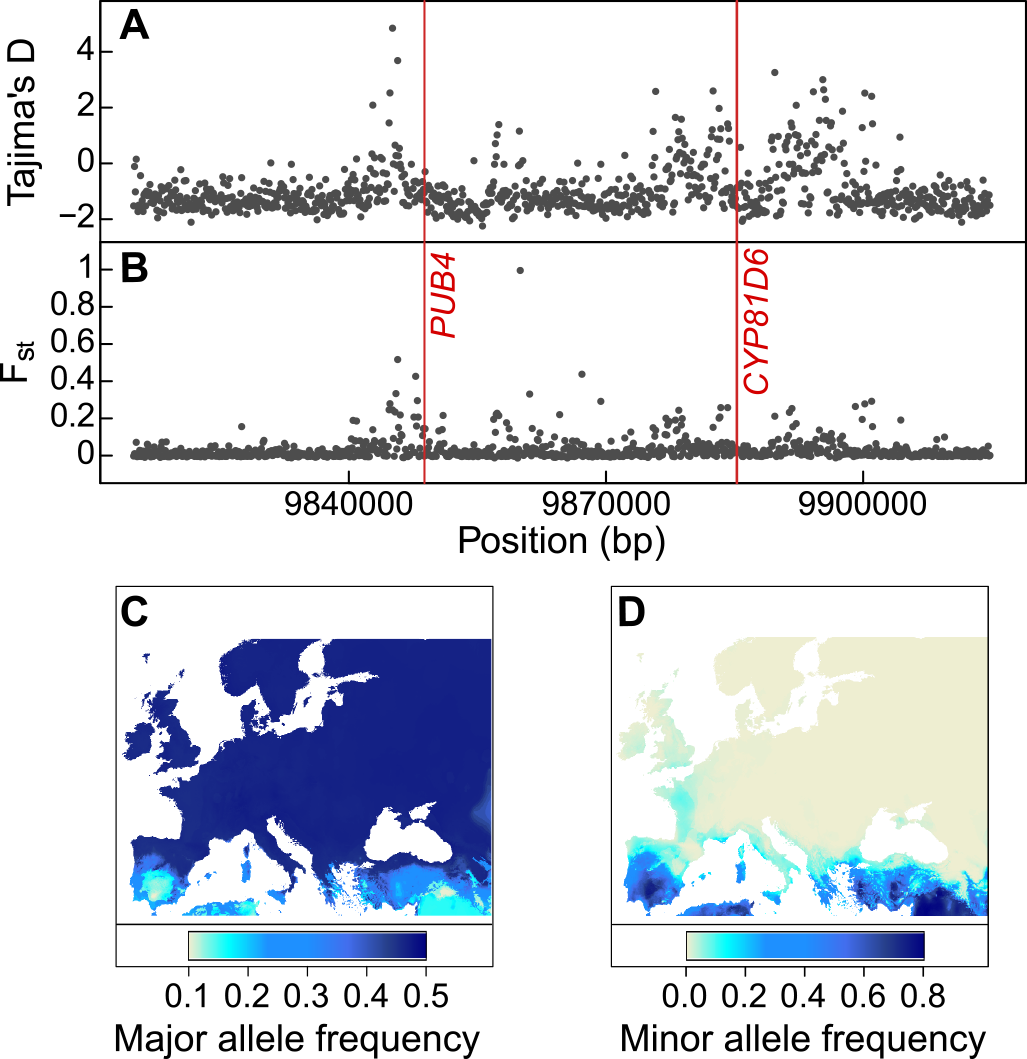
Genomic signatures of adaptation to climatic conditions at genes controlling the scaling exponent. (**A**, **B**) Tajima’s D (**A**) and F_st_ (**B**) in a 50 kb region around *PUB4* and *CYP81D6*. Grey dots are mean values in 1 kb-bins, red lines indicate positions of significant SNPs. (**C**, **D**) Predicted geographic frequency of the major (**C**) and minor (**D**) alleles at *PUB4* following climate-envelope modelling with 19 Bioclim variables. Color gradient indicates predicted allele frequency.

## Discussion

Metabolic allometry links physiology, ecology and evolution at different levels of organization (4, 6, 37, 38). The study of scaling relationships in both plants and animals is grounded on the importance of universal metabolic properties that allow the measurement and prediction of critical rates of energy flow from individuals to the biosphere (6, 39). However, explanations for the origin of allometric variation between species remain elusive, despite a recognized role of evolutionary processes in animals (40). Changes in scaling intercept in response to selection are well documented (41), but evidence for the evolution of allometric slopes is scarce (but see (42)), in particular in plants where the focus has been on the specific value that the allometric slope should take (*e.g.* ⅔ versus ¾ versus 1) (9, 13, 43).

Our results reconcile recent debates on the origin of biological allometry. On the one hand, our results support the idea that growth allometry varies significantly and that genetic variation in allometry is maintained within species. On the other hand, the canonical ¾ scaling exponent reported within and across plant and animal species was found to be associated with a phenotypic optimum that maximizes fitness under benign conditions, consistent with a role of stabilizing selection (4). Nonetheless, depending on the local environment, deviations in both directions from the ¾ scaling exponent might be advantageous for stress resistance despite their cost on seed production. Thus, stabilizing selection on metabolic allometry could be disruptive under unfavorable environments, as we have found for *A. thaliana*. Allometric adaptation may be due to, for instance, selection for fast growth and short lifespan to escape drought, or selection for resistance to hydraulic cavitation associated with reduced stomatal conductance and carbon assimilation in late flowering ecotypes (23, 44).

Specifically, these findings shed light on the important role of allometry for local adaptation to various climates in *A. thaliana*. Moreover, our results inform our understanding of the evolutionary basis of the tenets of MST. The maintenance of high intermediate-frequency nucleotide diversity in genes affecting allometry could result from long-term, geographically heterogeneous selection to optimize growth and survival in contrasting environments. This appears to have resulted in the genetic diversification of the scaling exponent around the intra- and interspecific mean of ¾, potentially reconciling the original MST prediction of an optimal scaling ¾ value with observed departures from it that have generated past debates (45). An intriguing question is whether the observed variation in scaling exponents across species (46) is associated with a similar climate adaptation as we observed for *A. thaliana*. Inter- and intraspecific variation in the vascular network and its impact on hydrodynamic resistance, resource distribution and plant allometry is already being explored (47, 48). If genetic variability in growth allometry is confirmed in other species and associated with climate, this would have important implications for our understanding of the physiological bases of plant adaptation. Moreover, it would connect macroevolutionary patterns of trait covariation observed across species to microevolutionary processes occurring within species.

## Materials and Methods

### Published data

For stress resistance analysis, we used published data from two studies on the response of *A. thaliana* natural accessions to drought: one where 210 accessions shared with our study were subjected to severe, lethal drought and survival was estimated for all accessions (21), and one where 60 shared accessions were subjected to 7 d non-lethal drought and fresh weight measured (22). We also re-analyzed phenotypic data previously published (10, 23) from a population of 120 L*er*-2 x Cvi recombinant inbred lines (49), and grown under water deficit and high temperature (10, 23).

Climatic data consisted of 19 bioclimatic variables (http://www.worldclim.org/bioclim) with a 2.5 arc-minutes resolution for the 1950 to 2000 CE period, plus mean annual Potential Evapo-Transpiration (PET, mm) and annual Aridity Index downloaded from http://www.cgiar-csi.org/data/global-aridity-and-pet-database (19). Monthly averages were calculated with 30 arc-seconds (ca. 1 km). Additional details in SI.

### Plant Material and Growth

We selected 451 natural accessions of *Arabidopsis thaliana* from the 1001 Genomes project (17) (http://1001genomes.org/; Dataset S1). Seeds were from parents propagated under similar conditions in the greenhouse. Four replicates of each accession were grown, with one replicate each sown on four consecutive days. Two replicates per accession were harvested as 16 day-old seedlings for dissection, imaging and weighing, and two were cultivated until the end of the life cycle (until fruit ripening) for trait measurement. Plants were cultivated in hydroponics culture on rockwool. Seedlings were vernalized for 4°C (8 h light) for 41 days. Plants were then transferred to 16 °C (12 h light). Additional details in SI.

### Plant Measurements

The Raspberry Pi Automated Plant Analysis (RAPA) system was used for continuous imaging using 192 micro-cameras (OmniVision OV5647), which simultaneously acquired 6 daily top-view 5 Megapixel images for each tray of 30 plants during the first 25 days after vernalization. Recording and storage of images were managed through embedded computers (Raspberry Pi rev. 1.2, Raspberry Pi Foundation, UK). Inflorescences and rosettes of mature plants were separated and photographed (Canon EOS-1, Canon Inc., Japan). The rosette was dried for at least three days at 65 °C, and weighed with a microbalance (XA52/2X, A. Rauch GmbH, Graz, Austria).

Fruits (siliques) were counted by eye on inflorescence images of 352 plants harvested at maturity. We analyzed the inflorescence pictures of all harvested plants with ImageJ (50) to estimate the number of fruits through image 2D skeletonization (18). The inferred variables were used to predict fruit number with linear regression (*glm*) performed on the 352 plants for which we had both measurements (18).

Drought survival index were from published data, measured as the quadratic coefficient of the polynomial regression between green leaves and time after the end of watering; more negative values mean lower survival (21). Measurements of growth reduction under moderate drought were also from published data, measured as the percentage of rosette fresh weight after seven days of water deficit compared to control (22). In the re-analysis of the population of 120 RILs previously phenotyped for growth scaling exponent (10), and trait plasticity in response to water deficit and high temperature (23), we measured resistance to combined stresses through the log ratio of dry mass under stress or no stress. Additional details in SI.

### Modeling Growth and RGR

Absolute growth rate (mg d^−1^) was estimated as the ratio of final rosette dry mass and plant lifespan. Using rosette dry mass estimated from image analysis (18), we fitted a sigmoid curve as a three-parameter logistic equation (51) with the function *nls* in R. From the parameters of the fitted function of each individual, we measured RGR (rosette growth rate divided by rosette dry mass, mg d^−1^ g^−1^) at the inflection point of the growth trajectory (18).

### Statistical Analyses

Statistical analyses except genomic analyses were performed in R (52). The coefficients of correlation (and their associated *P*-values) reported between phenotypic traits and climatic variables were the Pearson’s product moment coefficients obtained with the function *cor.test* in R. Effect of population structure on the scaling exponent was tested with ANOVA, using the nine genetic groups identified in the 1001 genomes dataset (http://1001genomes.github.io/admixture-map/) after removing admixed accessions (17). Broad-sense heritability (*H*^2^) was measured as the proportion of variance explained by genotype (Vg) over total variance (Vg + Ve) in a linear mixed model fitted with the ‘lme4’ R package, such as: *H*^2^ = Vg/(Vg + Ve). Similarly, Q_st_ was measured as the amount of variance in phenotypes explained by genetic group membership. As for *H*^2^, we used linear mixed model in the package ‘lme4’ in R to fit traits against genetic groups (nine genetic groups after removing ‘admixed’ accessions).

### Genetic Analyses

Conventional genome-wide association (GWA) studies were performed with easyGWAS (25) (https://easygwas.ethz.ch/). We used 1,793,606 SNPs with a minor allele frequency (MAF) above 0.05 to compute the realized relationship kernel from the full sequence of the accessions (http://1001genomes.org/). Association analyses were performed with EMMAX (53). For polygenic GWA, we used the Bayesian Sparse Linear Mixed model (BSLMM) implemented in GEMMA (30). Gene Ontology (GO) analysis was performed online using AgriGO (http://bioinfo.cau.edu.cn/agriGO/) (31) and REVIGO (http://revigo.irb.hr/) (54).

Prior to F_st_ calculation, genetic groups in the 1001 Genomes collection had been defined by ADMIXTURE clustering (55) (http://1001genomes.github.io/admixture-map/) (17). Genome-wide estimates of Weir and Cockerham F_st_ (56) were obtained with PLINK v1.9 (57). Local selection scans (Tajima’s D and F_st_) were obtained in 1 kb sliding windows in the 50 kb region around *PUB4* using PLINK. Selection sweep scans were carried out using SweeD software (58). Additional details in SI.

### Modeling Geographic Distribution

We performed stepwise regression to identify the set of climatic variables that best explain the variation of the scaling exponent between 36°N and 64°N, and 10.5°W and 27.5°E. We then used linear regression of the scaling exponent with the 13 best climatic variables to predict the exponent at every location across Europe. Geographic representation was obtained with the package ‘raster’ in R. We performed climate-envelope modelling of allelic frequency at *PUB4* with *maxent* modelling (59), using the package ‘dismo’ and ‘raster’ in R. We used the 19 Bioclim variables downloaded from Worldclim database at the origin of accessions with a 2.5 arc-minutes resolution. Additional details in SI.

## Acknowledgments

We thank Rebecca Schwab and Justine Bresson for their help in preparing and performing the experiments. We thank members of the Weigel and Burbano labs for their comments on previous versions of the manuscript. This work was supported by the sabbatical fellowship program from Colegio de Postgraduados de Montecillo (OJAG), an NSF ATB and Macrosystems award and a CNRS Associate Research Fellowship (BJE), ERC (ERC-StG-CONSTRAINTS, CV; ERC-AdG-IMMUNEMESIS, DW), and the Max Planck Society (FV, MEA, GW, DW).

## Author Contributions

FV, CV and DW designed the study. FV performed the experiments and extracted the data. FV, OJAG, DV, GW and MEA performed statistical analyses. All authors interpreted the results and wrote the paper.

## Data Availability

Phenotypic data are available in SI and on Dryad repository (http://datadryad.org/). R codes and ImageJ macro for data analysis are available on Github (https://github.com/fvasseur). GWAS results are available in easyGWAS (https://easygwas.ethz.ch/).

